# Predatory Behavior is Primary Predictor of Movement of Wildland-Urban Cougars

**DOI:** 10.1101/257295

**Authors:** Frances E. Buderman, Mevin B. Hooten, Mat W. Alldredge, Ephraim M. Hanks, Jacob S. Ivan

**Affiliations:** Colorado State University, 1484 Campus Delivery, Fort Collins, Fort Collins, CO 80523-1484; U.S. Geological Survey, Colorado Cooperative Fish and Wildlife Research Unit, Colorado State University, 1484 Campus Delivery, Fort Collins, CO 80523, USA; Colorado Parks and Wildlife, 317 W Prospect Road, Fort Collins, CO 80526, USA; Pennsylvania State University, W-250 Millennium Science Complex University Park, PA 16802

**Keywords:** cougar, individual variation, movement, population-level, predation, wildland- urban interface

## Abstract

While many species have suffered from the detrimental impacts of increasing human population growth, some species, such as cougars (*Puma concolor*), have been observed using human-modified landscapes. However, human-modified habitat can be a source of both increased risk and increased food availability, particularly for large carnivores. Assessing preferential use of the landscape is important for managing wildlife and can be particularly useful in transitional habitats, such as at the wildland-urban interface. Preferential use is often evaluated using resource selection functions (RSFs), but RSFs do not adequately account for the habitat available to an individual at a given time and may mask conflict or avoidance behavior. Contemporary approaches to incorporate landscape availability into the assessment of habitat preference include spatio-temporal point process models, step-selection functions, and continuous-time Markov chain (CTMC) models; in contrast with the other methods, the CTMC model allows for explicit inference on animal movement. We used the CTMC framework to model speed and directionality of movement by a population of cougars inhabiting the Front Range of Colorado, U.S.A., an area exhibiting rapid population growth and increased recreational use, as a function of individual variation and time-varying responses to landscape covariates. The time-varying framework allowed us to detect temporal variability that would be masked in a generalized linear model. We found evidence for individual- and daily temporal-variability in cougar response to landscape characteristics. Distance to nearest kill site emerged as the most important driver of movement at a population-level. We also detected seasonal differences in average response to elevation, heat loading, and distance to roads. Motility was also a function of amount of development, with cougars moving faster in developed areas than in undeveloped areas.

---

Quantifying individual variability in habitat selection, while simultaneously estimating population-level patterns, can be important for management and conservation issues where resources are heterogeneous or cause points of conflict (Kertson et al. 2011). Some large carnivores, such as cougars (*Puma concolor*), have undergone recent range expansions into human-modified landscapes (Knopff et al. 2014), but they rarely use the heavily modified landscapes in urban and suburban areas, instead relying on the rural and exurban areas at the wildland-urban interface (Burdett et al. 2010; Kertson et al. 2011). Along with increased risk from human interactions (Burdett et al. 2010), human-modified landscapes may contain greater numbers of both primary (ungulates, e.g., Torres et al. 2011) and secondary (domestic animals, e.g., Torres et al. 1996) prey for large carnivores compared to adjacent wild-land areas.

As early as 1998, the frequency of human-cougar interactions along portions of the Front Range, a mountain range extending north-south from Casper, Wyoming to Pueblo, Colorado, had increased due to encroaching residential development, increasing cougar populations, and increasing prey densities near human populations (Manfredo et al. 1998). The Front Range Urban Corridor runs along the eastern edge of the Front Range, while the Front Range itself contains a matrix of towns and areas that are managed for recreational use by county, state, and federal agencies. Human-cougar interactions have remained high in recent years (Mat Alldredge, Colorado Parks and Wildlife, personal communication), and cougars have been observed using developed areas in the Front Range as a reliable hunting ground (Blecha 2015; Moss et al. 2016). In addition, due to their desirable qualities, regions adjacent to protected areas have demonstrated higher population growth compared to growth in rural, non-protected areas (Wittemyer et al. 2008), increasing the potential for human-wildlife conflict (White and Ward 2011).

Individual-level movement decisions are one of the underlying processes that give rise to population-level patterns such as species distributions or their density and abundance on the landscape (Wiens et al. 1993). Movement decisions are a function of a number of variables, including the current location of the individual and the alternative available landscape (Wiens et al. 1993). Therefore, a central theme of animal ecology is the assessment of an individual’s preference for habitat, given what is available (Johnson 1980). Habitat preference is typically characterized using resource selection functions (RSF), which are often fit using logistic regression to compare the locations used by an individual or population to a random sample taken across some area defined as “available” (Manly et al. 2007). Use that is disproportionate to habitat availability implies that the individual has a preference for, or aversion to, the given habitat (Manly et al. 2007). However, inference on preference depends on what components are considered available to the animal (Johnson 1980). For example, an animal may use a resource disproportionately less than is available in its home range, however it may have chosen its home range because the resource was abundant (Johnson 1980).

In addition, availability is constrained by an individual’s range of movement. To account for dynamic availability, spatio-temporal point process models simultaneously estimate the resource selection function and time-varying availability kernels, which is the area an individual is capable of moving to over a given period of time (Christ et al. 2008; Johnson et al. 2008b; Brost et al. 2015). The more commonly used method, a step-selection function, approximates the availability kernel by using conditional logistic regression and a sample of “available” steps that an individual could have taken (e.g., Boyce et al. 2003; Fortin et al. 2005; Forester et al. 2009). Recent methods have used conditional logistic regression to separately approximate the movement and time-varying availability kernels, in the vein of spatio-temporal point process models (e.g., Avgar et al. 2016). However, because all of these methods are formulated in discrete time, inference is made only when data were observed and not on the unobserved path. In addition, aside from the spatio-temporal point process of Brost et al. (2015), none of these methods account for measurement error.

In contrast to many resource selection studies, one of the primary goals of continuous-time movement models is to estimate the true path of an individual when it was unobserved (Johnson et al. 2008a; Patterson et al. 2008; Brost et al. 2015; Buderman et al. 2016; Hooten and Johnson 2017). Continuous-time movement models can also incorporate measurement error and irregular observations in time. However, movement models are typically time consuming and computationally intensive to fit, making it difficult to obtain inference on multiple individuals (Hooten et al. 2016). If inference on multiple individuals is attainable, it may be possible to identify a population-level response that is consistent across individuals, which would provide a rigorous link between individual choices and population-level patterns (Wiens et al. 1993). In addition, understanding individual variability may help identify individuals that associate more strongly with certain features of the landscape (Aune 1991).

A recently developed method, continuous-time Markov chain (CTMC) modeling, incorporates an explicit movement model to obtain information on travel speeds. Travel speeds provide indirect inference on resource selection (Dickson et al. 2005) and avoid absolute statements about preference (Johnson 1980). The CTMC method (Hooten et al. 2010; Hanks et al. 2015), is fit in two stages, where the first stage uses a continuous-time movement model to obtain inference on where the individual was when it was unobserved, while the second stage allows for evaluation of landscape drivers of animal movement. The second stage of the analysis uses a Poisson likelihood with an offset to model transition rates; therefore, statistical software based on a Poisson likelihood can implement the CTMC movement model (Hanks et al. 2015). The flexibility of the CTMC framework can account for time-varying responses to landscape drivers by allowing coefficients to vary temporally (Hanks et al. 2015), and it can also be implemented in a Bayesian hierarchical framework, allowing for inference on individual- and population-level drivers.

Given the increasing potential for human-wildlife conflict as development permeates rural and wildland areas along the Front Range and elsewhere in the West, we sought to extend previous work by explicitly modeling cougar movement to identify key drivers of their behavior, and in doing so, better understand their use of the wildland-urban landscape in both space and time. Many cougar studies do not explicitly model movement, and instead focus on resource selection, making inference from approximately one or fewer locations per day; these locations were sometimes obtained only during daylight (e.g., Anderson et al. 1992; Beier 1995; Dickson and Beier 2002), obtained during night and day but were treated equivalently (e.g., Hemker et al. 1984), or obtained at unspecified times (e.g., Ruth et al. 1998; Sweanor et al. 2000). Inference on time-varying behavior has been limited to separate analyses on discretized temporal periods (e.g., Dickson et al. 2005; Knopff et al. 2014). Some studies have also focused exclusively on kill site and hunting locations (e.g., Blecha 2015) or non-kill site locations (e.g., Dickson et al. 2005; Knopff et al. 2014). In contrast to previous studies, we used the CTMC framework to model individual- and population-level cougar responses to landscape features in continuous time, which allowed for direct inference on how behavior varied at a fine temporal scale, given what was available. In addition, by using a hierarchical modeling framework, we accounted for individual-level variation, which may be a function of the spatial distribution of prey items or the behavioral flexibility of a generalist predator (Kertson et al. 2011; Wilmers et al. 2013), while still obtaining population-level inference across a suite of individuals.

## METHODS

### Data Collection and Study Area

As part of an ongoing study by Colorado Parks and Wildlife (CPW), cougars were trapped and fit with global positioning system (GPS) collars and released along the Front Range of Colorado (CPW ACUC 01-2008; Figure 1). We were interested in examining seasonal differences in cougar movement due to temporal variability in the landscape-level covariates. For example, we expected a stronger response to prey-based covariates in summer months, because mule deer fawns, a primary prey source for cougars, are born in June (Pojar and Bowden 2004) and fawns are at a disproportionately high risk for predation (Hornocker 1970). We focused on 19 adult individuals (M=5, F=14) that were monitored April 1-15 2011, 21 adult individuals (M=7, F=14) that were monitored during June 16-30 2011, and 21 adult individuals (M=3, F=18) that were monitored October 1-15 2011. All individuals were monitored with Vectronics collars (Vectronics GmbH, Berlin, Germany) programmed to obtain fixes every three hours.

**Figure 1:**
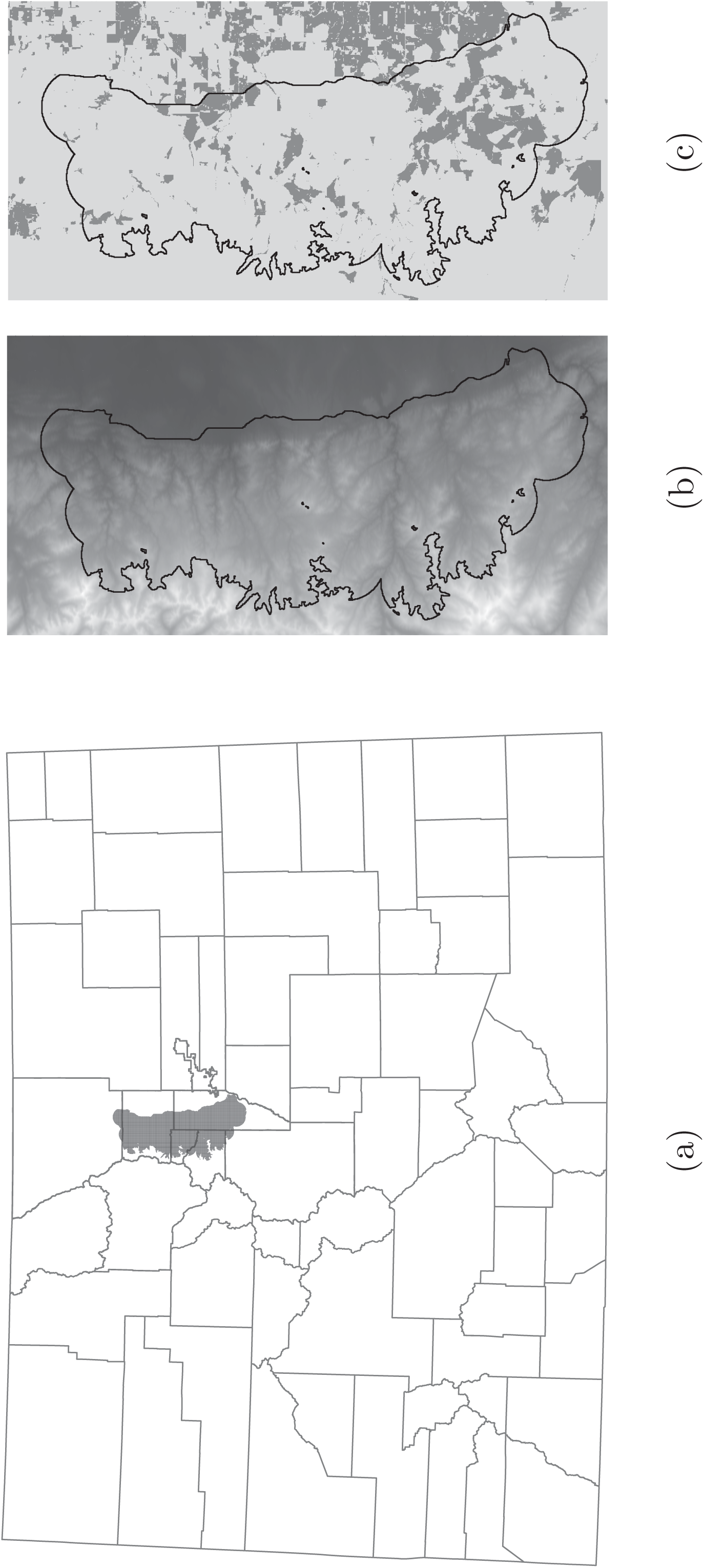
Map of Colorado counties, with the cougar movement study area plotted in gray (Figure 1a). Elevation (m; Figure 1b) and land classified as developed (dark gray is <10 acres/unit; Figure 1c) is shown for the study area and surrounding area.

Our study area comprised a 2,700 km^2^ region in the Colorado Front Range to the north- west of Denver (Figure 1). The study area consisted of a matrix of private (43%) and public (57%) land (Blecha 2015). Private land included areas of rural, exurban, suburban development, and small towns. Public land was managed by federal, state, and municipal governments for recreational activities or as open space. Road density was, on average 1.6 km/km^2^, but ranged from no roads to 16.6 km/km^2^. Elevation ranged between 1,522 and 4,328 m, generally increasing east to west, with development decreasing along a similar gradient. Ponderosa pine, Douglas fir, lodgepole pine, and spruce-fir were the predominant non-agriculture vegetation types, in order of dominance from east to west. Cougar density was approximately 2.4 individuals per 100 km^2^ (Mat Alldredge, CPW, personal communication).

### Stage 1: Continuous-Time Markov Chain Model

We used a Bayesian hierarchical CTMC model to evaluate drivers of cougar movement; this model is an extension of the model proposed by Hanks et al. (2015) and allows for inference on movement rates and directional bias, as opposed to use, in continuous time and discrete space. The initial step in the CTMC framework is to estimate a continuous movement path from the observed data points. We used the functional movement model proposed by Buderman et al. (2016) to account for observation error and predict locations every ten minutes for the selected two weeks of each month. A random subset of paths from the posterior predictive distribution of the movement model were spatially discretized to a latent variable formulation with a cell size of 100 m^2^, which was the largest cell size among the available covariates.

The CTMC model consists of a product of two components: the time an individual spends in a grid cell and the direction that an individual moves when it leaves a grid cell. The time an individual spends in a grid cell (motility) is exponentially distributed, such that a large rate parameter corresponds to fast movement out of the cell. When an individual leaves a grid cell, the probability that they move to a particular neighboring grid cell (directionality) is the ratio between the movement rate into that cell and the movement rate out of the preceding cell. Therefore, higher proportional rates indicate directional bias in movement. Thus, movement rate parameters, which are a function of covariates (i.e., landscape variables that correspond to the position of the cell on the landscape), control both motility and directionality. Hanks et al. (2015) showed that the likelihood of the CTMC model (i.e., the product of the motility and directional components) for movement can be expressed as a Poisson GLM using a latent variable formulation.

In the latent variable formulation, each transition corresponds to four data points (the four neighboring grid cells); the response variable is equal to one if the neighboring grid cell is the cell that the individual transitioned into and zero otherwise. Modeling the latent variables (zeros and ones) as Poisson random variables with an offset for the amount of time an individual spends in a grid cell results in a likelihood that is equivalent to the CTMC likelihood. This allows inference to be obtained using standard GLM software, and the R package `ctmcmove` facilitates creation of the CTMC latent variable formulation (Hanks 2017). Full CTMC details are available in Appendix A.

Using multiple imputed paths accounts for the uncertainty in the true path of the individual and is a process version of multiple imputation (Hooten et al. 2010, Hanks et al. 2015; Hooten et al. 2016; Scharf et al. 2017), a method frequently used for missing data (Rubin 1987). Process imputation is more computationally efficient than using the entire posterior distribution, but still approximates the uncertainty associated with the unobserved path. We generated 30 imputations for each individual, using 20 imputations to fit the models for individual-and population-level inference on transition rates and 10 imputations to calculate the posterior predictive score that was used to select regularization terms. Regularization shrinks the effect of unimportant covariates toward zero to prevent over-fitting (Hooten and Hobbs 2015).

### Stage 2: Poisson Models for Movement Inference

We assessed cougar response, as measured by movement rates and directional bias, to landscape features, including measures of anthropogenic activity. Because cougars and humans are active at different times throughout the day, we proposed two models for drivers of cougar movement: a hierarchical generalized linear model (H-GLM) for individual- and population-level inference on average cougar behavior and a hierarchical (e.g., mixed) generalized additive model (H- GAM) to account for individual- and population-level diel time-varying behavior. Covariates were centered and scaled to the individual, meaning that the coefficients are relative to the mean and standard deviation of the values that each individual encountered during a given two-week period. This is similar to the idea proposed by Johnson (1980), where preference was determined by comparing some measure of usage and availability of a landscape feature on an individual basis.

On average, we expected cougars to respond similarly to landscape covariates. Therefore, we developed a H-GLM for the latent variable formulation of the CTMC framework. In the CTMC framework, the response variables, *z*_*ij*_, are a sequence of zeros and ones, where *z*_*ij*_ ∼ Poisson(λ_*ij*_), for *i* = 1, …, *T* and *j* = 1, …, J, where *T* is the total number of cell transitions, and *J* is the number of individuals. Landscape covariates are incorporated using the log link function, such that 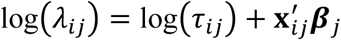. The residence times are represented by the constants ***τ*** _*ij*_, and the landscape variables by **x**_*ij*_. The parameter ***β*** _*j*_ is a vector of Pindividual-level coefficients that arise from the population-level distribution ***β***_*j*_∼ 𝒩(**μ**_*β*_, Σ_*β*_).The covariance matrix,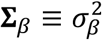 diag(***ϕ***), where the vector ***ϕ*** scales the value 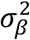 to each coefficient. The vector of scaling parameters consists of a one (***ϕ*** _1_= 1) and is modeled as log(***ϕ*** _p_) ∼N(0, 0.04) for p = 2, …, P. The population-level distribution has a mean that is modeled with a multivariate normal distribution 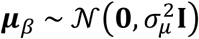. Both 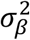and 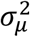are used as regularization terms, where 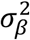 is selected *a priori* and 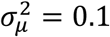, to shrink the coefficients toward zero; this prevents over-fitting and allows for correlated predictors (Hooten and Hobbs 2015).

The H-GAM is formulated as a varying coefficient model (Hastie and Tibshirani 1993), where the response to covariates varies over space or time. By expanding the landscape covariates with a basis function (Hefley et al. 2017), we created a new vector, **v**_*ij*_, that is the Kronecker product of the *P* length vector of covariates, 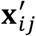, and the *Q* length vector of the values of the basis at the time of transition *i*, **w**(*i*). For diel movement, we used cubic cyclic spline basis functions (**w**(*i*)), because they constrain the start and end points of the varying coefficients to be equal, which is an important property for time spans that are cyclic in nature. The GAM for hourly movement is similar to the GLM, except 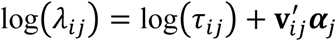, where ***α*** _*j*_ is a vector of length *PQ*. Each parameter in ***α*** _*j*_is the collective effect of the basis function and the corresponding covariate at the time of transition i. Using the vector w(i), ***α*** _*j*_ can be back-transformed to obtain the time-varying effect of the covariate. In the hierarchical framework ***α*** _*j*_∼N(μ_***α***_,Σ_***α***_),where 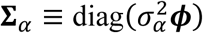.The vector ***ϕ*** again reduces the number of parameters we need to select *a priori* by scaling the 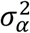 term to each parameter, and 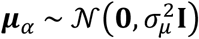. Both 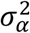 and 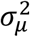 serve as regularization terms, where 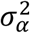 is selected *a priori* and 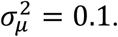.

Finally, to assess whether males and females exhibited different amounts of temporal variation in their response to potential movement drivers, we fit the GLM and GAM models to males and females separately for each time period. This resulted in four models: 1.) a GLM fit to all individuals, 2.) a GAM fit to all individuals, 3.) a GLM fit to females and a GAM fit to males, 4.) and a GAM fit to females and a GLM fit to males. We calculated the posterior predictive score for each model (i.e., the sum of the posterior predictive score for the models fit to males and females separately) and compared the scores across models within each month.

Models were fit using a Markov Chain Monte Carlo (MCMC) algorithm written in R (R Core Team 2016). We performed adaptive tuning over an initial 50,000 MCMC iterations. We used the selected tuning parameters as constants in the subsequent 50,000 iterations that were used to calculate the posterior predictive score for the *a priori* regularization parameter grid-search. The final models were fit using 100,000 MCMC iterations with a burn-in period of 10,000 iterations.

### Landscape Covariates

Each covariate can be included as either a motility or directional driver of movement in the CTMC model. Motility covariates are based on the value of the grid cell that the individual is in currently and control the absolute rate of movement; positive coefficients indicate faster movement with increasing values of the covariate (and slower movement with decreasing values), and negative coefficients correspond to faster movements with decreasing values of the covariate (and slower movement with increasing values). Directional covariates account for the correlation between movement and the gradient of a covariate and contribute to the probability that an individual moves toward a grid cell. The directional drivers were calculated such that a positive coefficient indicates that individuals move predominantly in the direction that the covariate decreases (decreasing distance, such that they orient toward a feature), whereas a negative coefficient indicates that individuals move in the direction that the covariate increases (increasing distance, orient away from a feature). All rasters were aggregated to a 100 m^2^ resolution, which is within the distance that a cougar might typically move over a ten-minute interval (Dickson et al. 2005).

We hypothesized that a number of landscape covariates may contribute to transition rates and directional bias of cougars: mule deer (*Odocoileus hemionus*) utilization (as a proxy for availability), distance to nearest potential kill site, distance to nearest structure, distance to nearest road, elevation, heat insolation load index, and topographic wetness. We also used an autoregressive parameter to account for an individual’s tendency to move in the direction it was already moving (directional persistence, Hanks et al. 2015).

Prey availability is a driving factor in cougar habitat selection. For example, cougars in western Washington used areas where suspected prey availability was high, such as low-elevation, early successional forests, and areas near water (Kertson et al. 2011), and Blecha (2015) observed cougars foraging in areas with high mule deer utilization. We approximated prey availability using two covariates: mule deer utilization and nearest potential kill site. The model averaged prediction for mule deer utilization (Blecha 2015) approximates prey availability given a suite of landscape covariates. We hypothesized that cougars would move slower in areas with high values for mule deer utilization and orient toward areas of high mule deer use during crepuscular and nocturnal movements (Anderson Jr and Lindzey 2003; Kertson et al. 2011; Blecha 2015). Blake and Gese (2016) found that many of the landscape variables that contribute to the location of predation events were the same as those contributing to non-predation habitat use, which led them to determine that cougars spend the majority of their time moving across the landscape in hunting mode. Including the location of a potential kill site may act as a proxy for unmeasured landscape variables and non-mule deer prey presence. Potential kill sites were determined using a clustering algorithm on the GPS points Knopff et al. (2009), where a location was classified as a potential kill site if two or more GPS locations, occurring between the average time of sunset and sunrise for each two-week period, were found within 200 m of the site within a six-day period. We calculated the distance (m) to nearest potential kill site across the study area to account for dependence in the movement process due to the known temporary activity centers induced by the potential kill sites. We expected individuals to move faster as distance to potential kill site increased, because decreasing distance may correspond to an individual returning to a cached kill, and caches are more often located in areas of high vegetation cover (Husseman et al. 2003).

We calculated distance to nearest structure (m) as the Euclidean distance to the nearest man-made roofed structure (Blecha 2015). Distance to road was calculated using major roads data (i.e., a major highway primarily for through traffic usually on a continuous route and streets whose primary purpose is to serve the internal traffic movement within an area) obtained from Colorado Department of Transportation. Due to increased human activity around structures and roads, we expected cougars to move faster when closer to roofed structure and distance to nearest road (Dickson and Beier 2002; Dickson et al. 2005; Nicholson et al. 2014). However, females may respond less to structures and roads than males, given that there may be additional factors, such as food limitation and offspring, which drive them to tolerate human-modified landscapes (Wilmers et al. 2013; Benson et al. 2016). We also expected there to be high temporal variability in the response to structures, because individuals have been observed avoiding areas of anthropogenic activity less at night, while avoiding contiguous forest habitat less during the day (Knopff et al. 2014).

We used a digital elevation model (Figure 1) to characterize elevation. Blecha (2015) found that cougars avoided foraging in higher elevations, but Wilmers et al. (2013) observed cougars selecting for higher elevations in developed areas. We expected cougars to show high temporal variability in their directional response to elevation, with cougars moving toward lower elevations when they are hunting (main prey is concentrated in lower elevations) and toward increasing elevations at other times. We used a raster based on the continuous heat insolation load index (CHILI, Theobald et al. 2015), modified from McCune and Keon (2002) to measure the accumulation of solar radiation at that location over the course of a year (MJ/cm^2^/yr). Heat insolation is high on south-facing slopes that are more xeric and open than north-facing slopes (Veblen and Donnegan 2005). Cougars have been observed using less rugged terrain for travel (Dickson et al. 2005), selecting for south-facing slopes containing shrubs (Knopff et al. 2014), and avoiding foraging on north-facing slopes (Blecha 2015). Therefore, we expected that cougars may orient toward areas of high heat insolation, but move quickly through them. The topographic wetness plus metric (TWI+) predicts soil moisture based on slope, as originally described by Beven and Kirkby (1979), and aspect, as modified by Theobald (2007). Because cougars have been observed selecting for and hunting in riparian areas (Dickson and Beier 2002; Kertson et al. 2011; Nicholson et al. 2014; Benson et al. 2016), we expected cougars to move slowly in areas of high topographic wetness and demonstrate temporal variability in their directional response (toward areas of increasing topographic wetness when hunting).

We also analyzed a subset of individuals and the interaction between housing density and their response to deer utilization and distance to nearest kill site. Despite cougars demonstrating avoidance of high housing densities while foraging, kill sites were positively related to housing density (Blecha 2015). In addition, the temporal variability in the response to anthropogenic structures that was observed by Knopff et al. (2014) was stronger for cougars in rural, rather than wilderness, areas. Therefore, these are the two variables that we expected to vary most with housing density due to the potential trade-offs between increased prey abundance but increased mortality risk. To determine the effect of housing density on the response of cougars to deer utilization and potential kill sites, we discretized the landscape into developed (<10 acres/unit) and undeveloped areas (Figure 1). Only 13, 15, and 17 individuals for April, June, and October, respectively, were used in the secondary analyses because the remaining individuals did not spend time in developed areas in the selected two-week periods. We were unable to evaluate an interaction in the H-GAM framework due to high variability in the percentage of locations for each individual that were classified as occurring in undeveloped areas.

## RESULTS

There was no detectable effect of many of the landscape covariates on average motility or directionality at a population-level (Figure 2). However, distance to potential kill site emerged as the primary driver of both motility and directionality in the GLM framework (Figures 2, 3). As individuals increased their distance from a potential kill site, their transition rate increased (Figure 3b). In addition, individuals oriented movement toward their potential kill site (Figure 3b). Although not significantly different from zero, the strength of the estimated effect of distance to kill site increased as the months progressed (Figure 2). We also detected significant residual autocorrelation, indicating that individuals tended to continue moving in the direction they had previously been moving, after accounting for landscape features (Figure 2).

**Figure 2:**
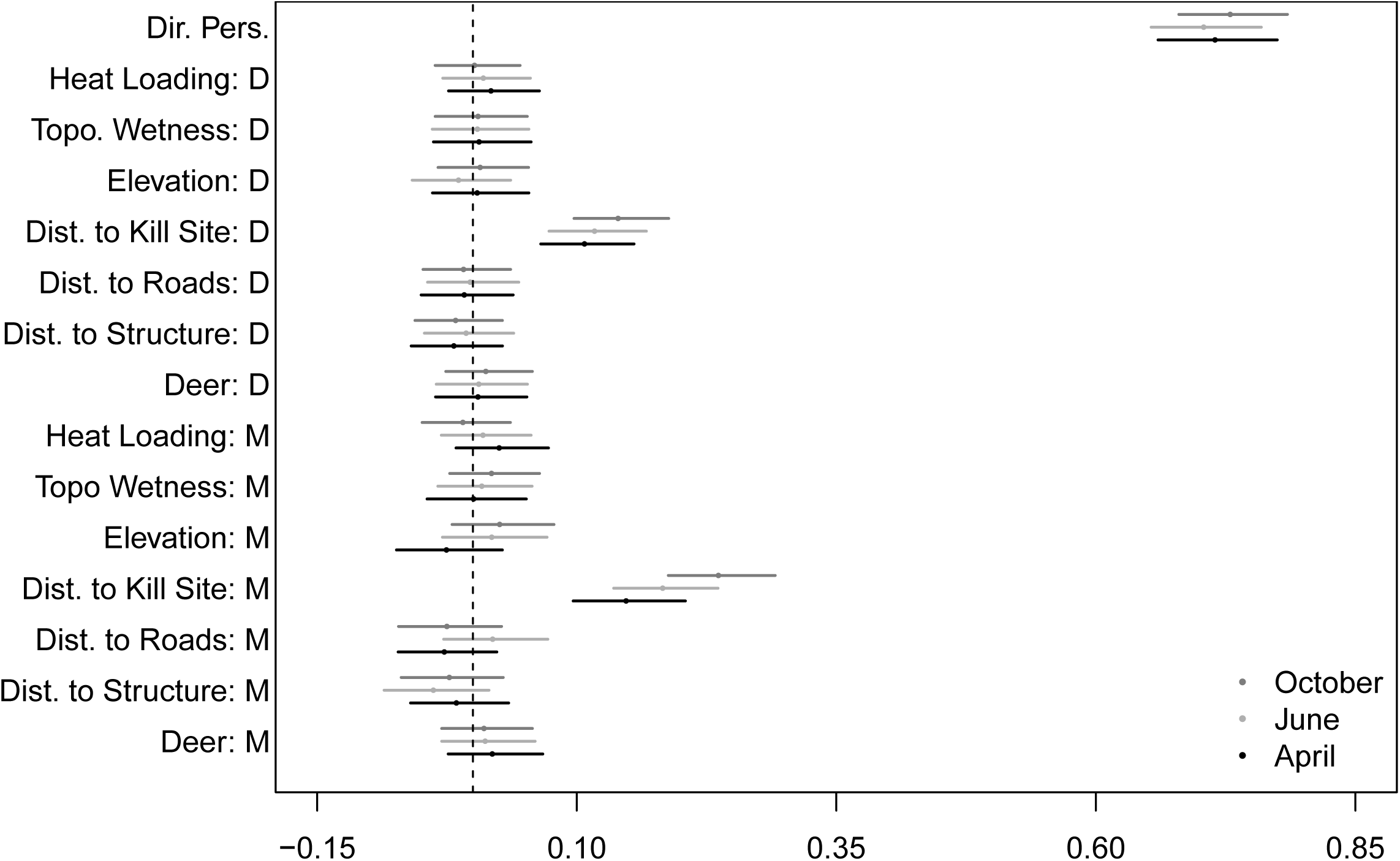
The mean and 95% credible intervals for the population-level mean effects of landscape covariates on movement rates (M) and directionality (D) of cougar movement in the Colorado Front Range for two-week periods in April, June, and October 2011.

**Figure 3:**
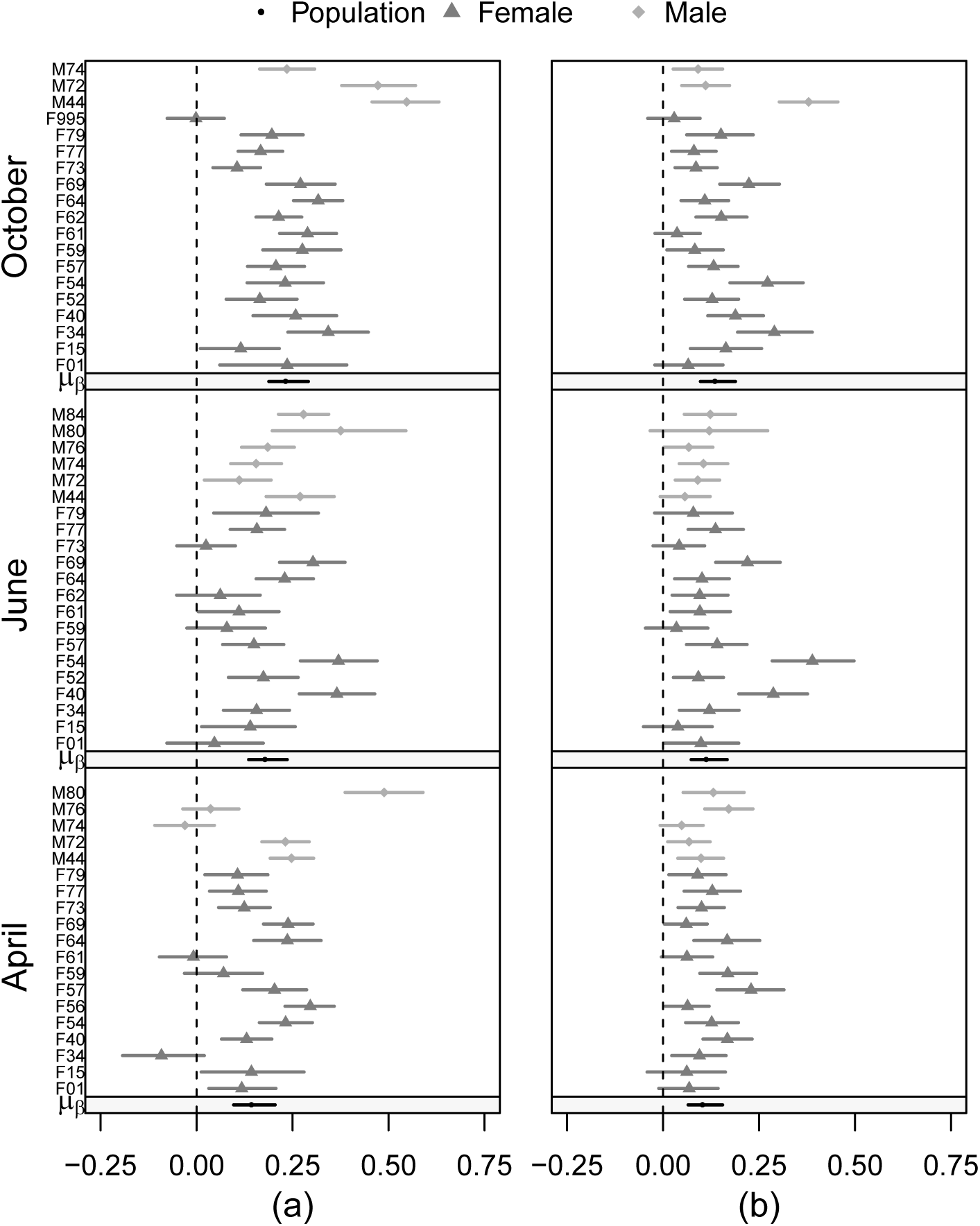
Posterior means and 95% credible intervals for the individual- and population-level static effects of distance to nearest potential kill site on motility (Figure 3a) and directionality (Figure 3b) of cougar movement in the Colorado Front Range for two-week periods in April, June, and October 2011.

Of the remaining potential drivers of movement, the largest seasonal differences were observed in the effect of heat loading, elevation, distance to nearest roofed structure, and distance to nearest road; however, the 95% credible intervals consistently overlapped zero (Figure 2). Based on the posterior mean, individuals were observed moving slower at higher elevations in April, but faster in June and October (Figure 4a). In contrast, individuals moved faster than average in areas where heat loading was high in April, and to a lesser degree, June, but moved slower with higher heat loading in October (Figure 4b). We detected a consistent seasonal effect of distance to structure, but the posterior mean was negative, meaning that individuals moved faster as distance to structure decreased (Figure 4c). A similar pattern was detected with distance to roads, however the effect became positive in June, with individuals moving faster as distance to roads increased (Figure 4d). In most cases, individual-level uncertainty tended to be high, with a few individuals showing statistically significant responses despite a non-significant population-level response (Figures 3, 4).

**Figure 4:**
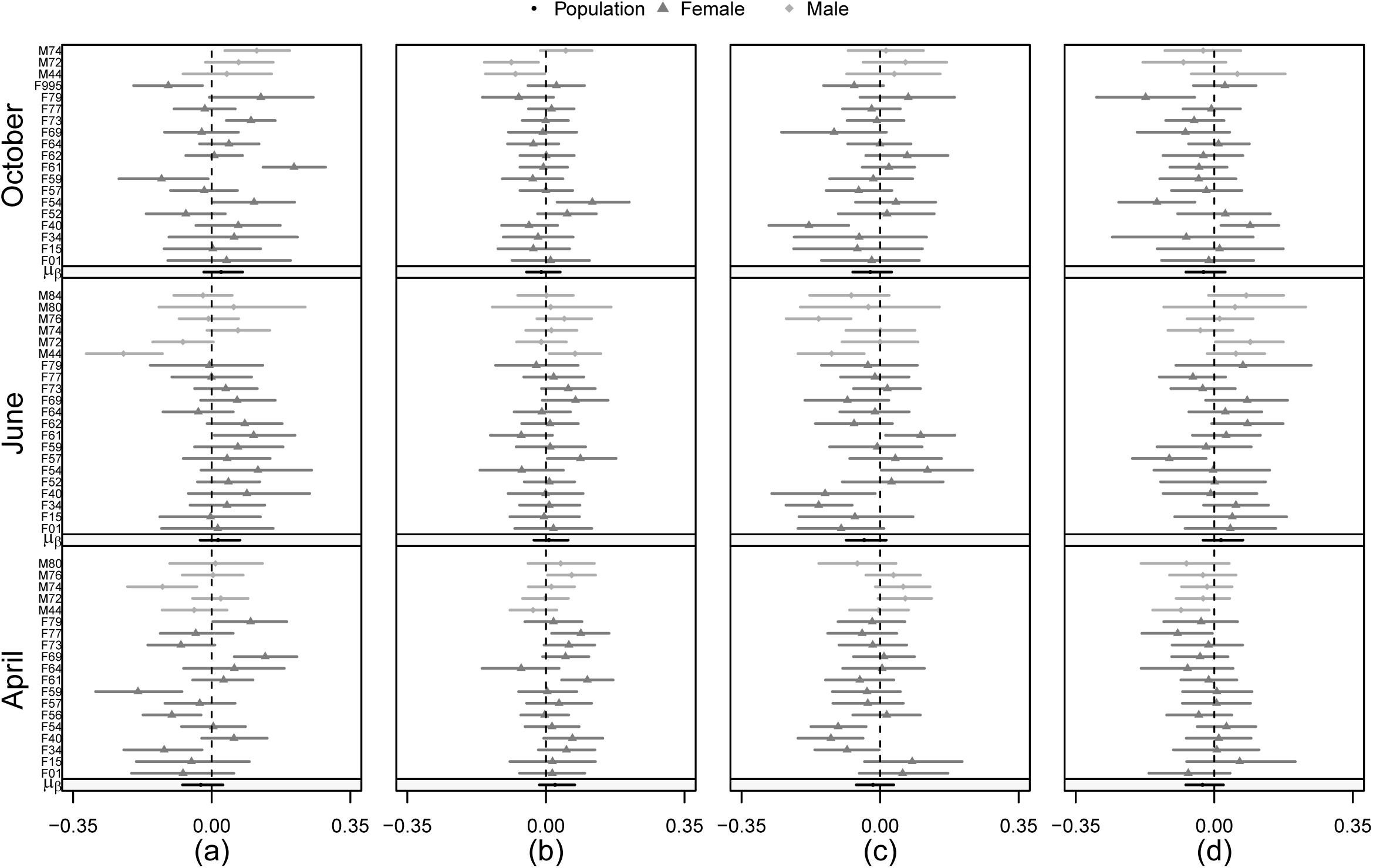
Posterior means and 95% credible intervals for the individual- and population-level static effects of elevation (Figure 4a), heat loading (Figure 4b), distance to nearest roofed structure (Figure 4c), and distance to nearest road (Figure 4d) on cougar motility in the Colorado Front Range for two-week periods in April, June, and October 2011.

Our results suggest that distance to nearest potential kill site was also the predominant motility and directionality driver in the diel time-varying framework (H-GAM; Figures 5 and 6). However, the strength of the motility response to distance to nearest potential kill site varied over time and with seasons. The strongest motility response occurred around dawn, decreased steadily during daylight hours, and then increased around dusk (Figure 5). The magnitude of this variation was strongest in June and October, and weakest in April (Figure 5). The strength of the directional bias toward potential kill sites also varied through time, but was consistent across seasons (Figure 6). The evidence for an effect of potential kill sites on directionality suggests that individuals orient less toward their kill site during daylight hours, and may even orient away from their potential kill sites during late afternoon (Figure 6).

**Figure 5:**
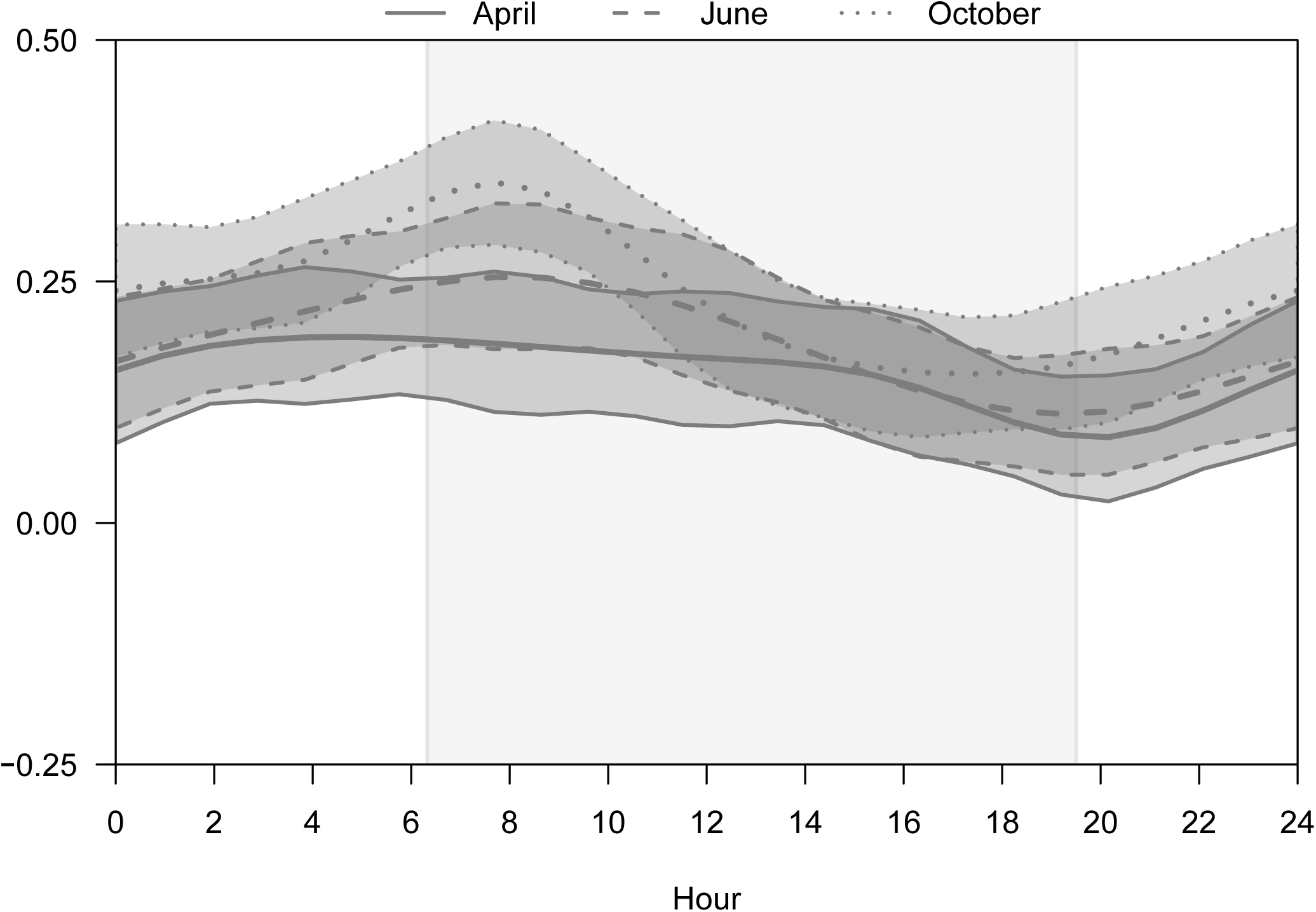
Posterior means and 95% credible intervals for the population-level diel time-varying effect of distance to nearest potential kill site on cougar motility in the Colorado Front Range for two-week periods in April, June, and October 2011. The gray box represents 0630 hours to 1930 hours.

**Figure 6:**
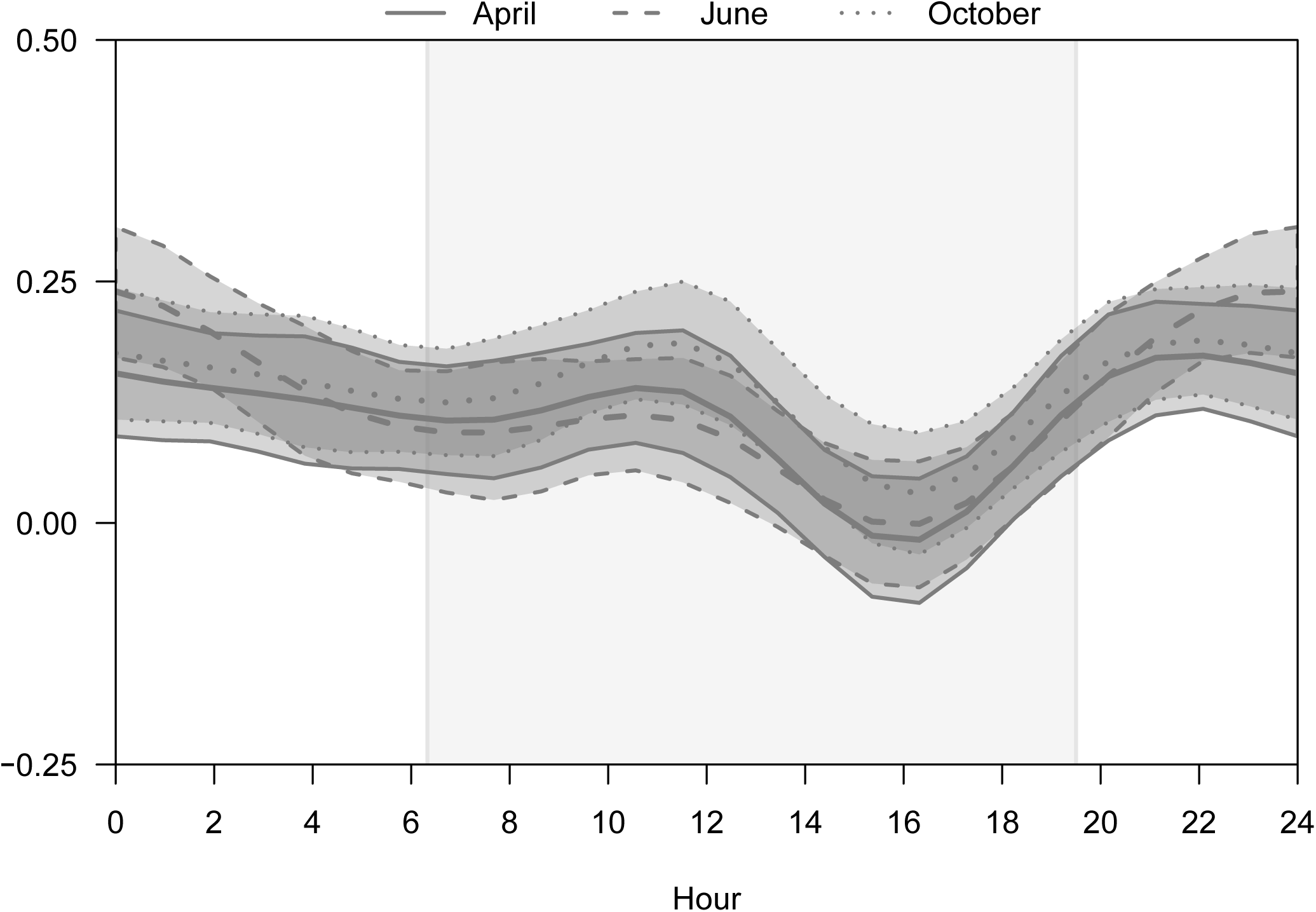
Posterior means and 95% credible intervals for the population-level diel time-varying effect of distance to nearest potential kill site on directionality of cougar movement in the Colorado Front Range for two-week periods in April, June, and October 2011. The gray box represents 0630 hours to 1930 hours.

While the 95% credible intervals overlapped zero for much of the day, we detected modest temporal responses in both motility and directionality to elevation and distance to nearest structure (Figure 7). The motility response to elevation varied seasonally, as in the GLM framework. The average negative response to elevation observed in April (Figure 4a) was reflected in a negative response to elevation around dawn (individuals move slower as elevation increases), with a slightly positive response later in the day (Figure 7a). We observed little time variation in June, but the pattern observed in October was the opposite of April, with individuals moving faster with increasing elevation around dawn, with a decreasing effect through the rest of the day (Figure 7a). Individuals moved toward higher elevations mid-day and toward lower elevations at other times, a pattern that was consistent across seasons (Figure 7b). The strongest negative effect of distance to structure on motility (individuals move faster as distance decreases) occurred around dawn and dusk for all seasons (Figure 7c). The effect on directionality was less consistent, with orientation toward roofed structures just after dawn, followed by orientation away from structures, in April and June; this pattern shifted toward pre-dawn in October (Figure 7d).

**Figure 7:**
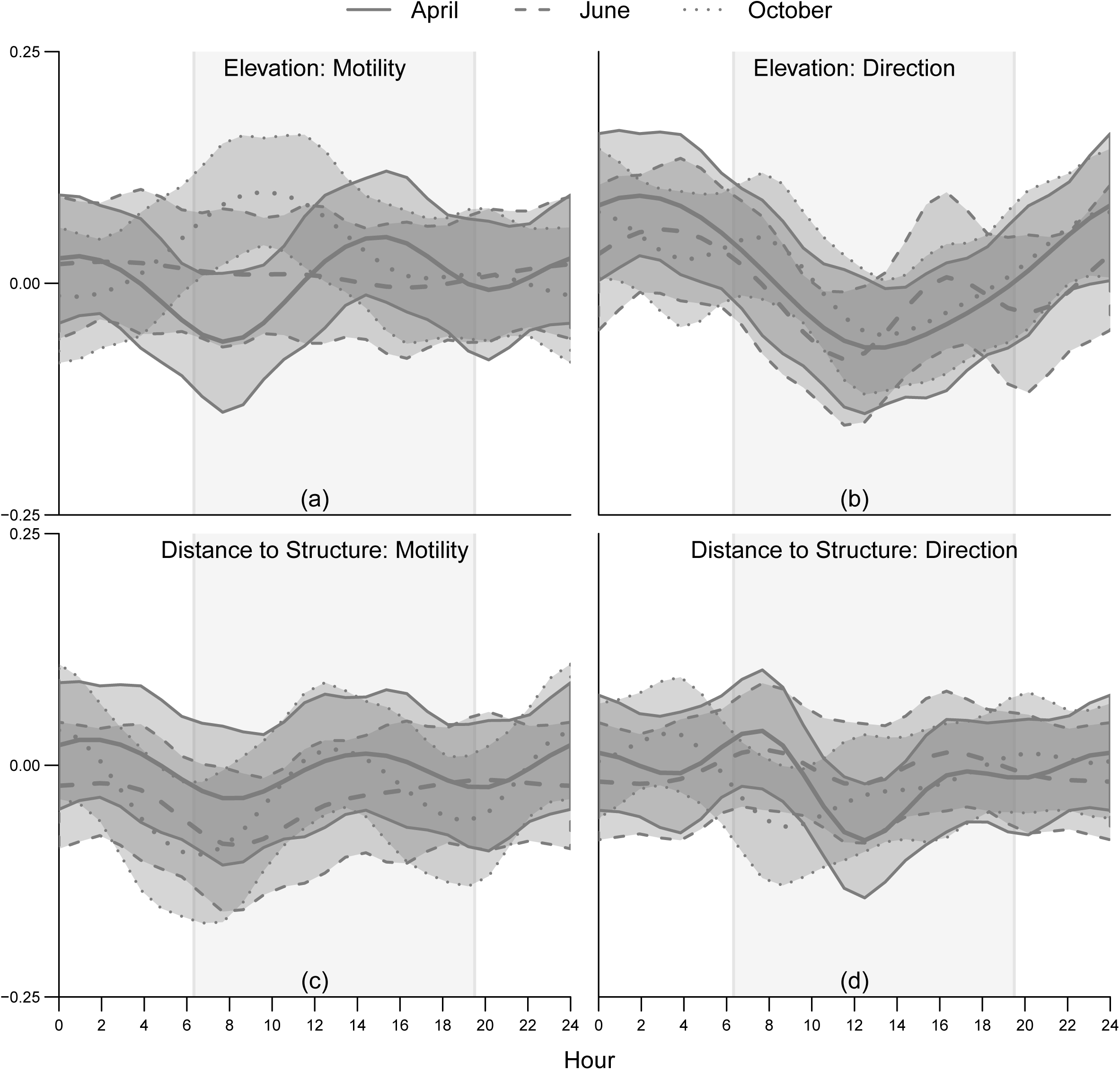
Posterior means and 95% credible intervals for the population-level diel time-varying effect of elevation (Figure 7a, 7b) and distance to nearest roofed structure (Figure 7c, 7d) on motility and directionality of cougar movement in the Colorado Front Range for two-week periods in April, June, and October 2011. The gray box represents 0630 hours to 1930 hours.

In addition, we did not see evidence for an interaction between development and deer utilization, which remained a statistically insignificant driver of cougar movement rates and directionality in both the H-GLM and H-GAM models. The positive effect of distance to potential kill site on speed (faster as distance to kill site increases) and directional bias (more orientation toward the kill site) was consistent between developed and undeveloped areas (Figure 8a). However, we detected a difference in average movement rate between the two areas, with individuals in each month moving faster in developed areas (Figure 8b).

**Figure 8:**
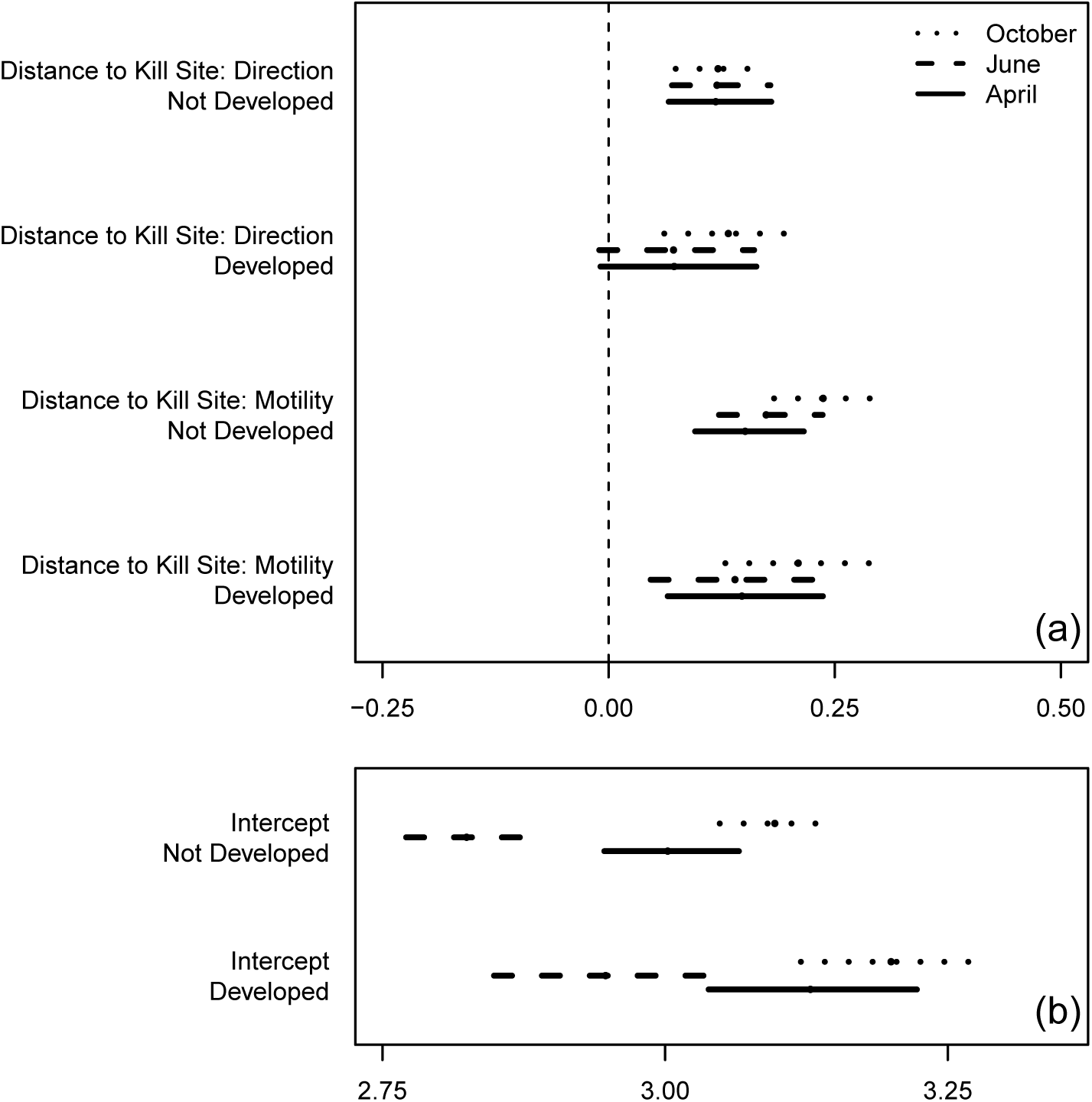
Posterior means and 95% credible intervals for the population-level effect of distance to nearest potential kill site on motility and directionality of cougar movement (Figure 7a) and average movement rate (Figure 7b) as a function of development (developed being <10 acres/unit) in the Colorado Front Range for two-week periods in April, June, and October 2011.

Finally, the H-GAM for both sexes was the best model in terms of predictive performance across all months, whereas the GLM performed the worst (Figure 9). The models that were a mixture of a GAM and GLM, varying by sex, were generally equivalent (Figure 9). The largest difference between the two sex-varying models was observed in October, when the better of the two models included time variation for males and no time variation for females (Figure 9).

**Figure 9:**
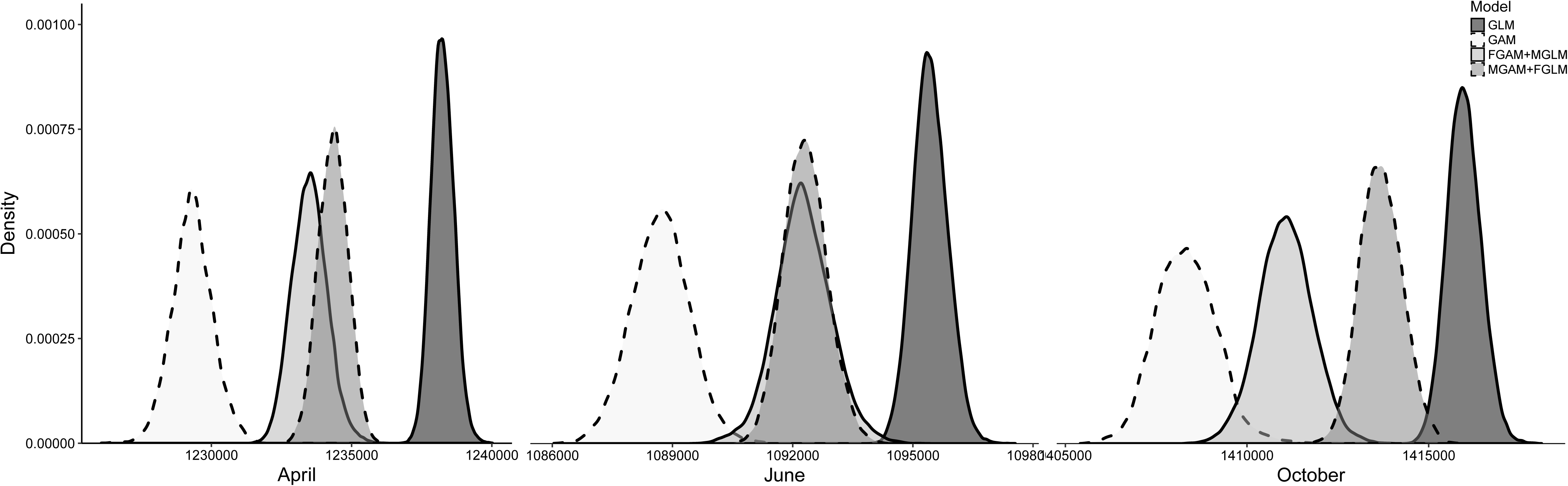
Posterior predictive score distributions for the model set, where smaller values indicate better predicting models. The GAM fit to all individuals performed the best in terms of predictive ability for all months.

## DISCUSSION

The strong observed response to distance to nearest potential kill site is likely due to returning visits to the carcass and unmeasured fine-scale covariates related to landscape features that increase the likelihood of a successful hunting attempt. Blecha (2015) found that kill sites, compared to preceding locations, occurred more frequently in areas with higher housing densities and lower topographic positions, such as drainage areas, despite drainage areas having lower prey availability. We did not measure hunting success, but we did find that cougars moved toward lower elevations at dusk, when cougars are likely to hunt or return to a carcass. We found that individual response to nearest potential kill site was variable within the two-week period and across individuals; this is likely a function of timing of successful kills and the size of the prey item, with stronger positive responses being correlated with larger prey. In addition, the variation in motility across months may also be a function of the available prey items during a given season. We observed the weakest response to kill sites during April, before mule deer fawns are born (Pojar and Bowden 2004) and when cougars are relying on smaller prey. We also observed an increasingly strong positive response to distance to kill site from dusk to dawn, implying that, from dusk to dawn, individuals moved increasingly faster the farther away they were from a potential kill site.

Our results indicated that cougars moved slightly faster in areas with a higher heat insolation load index in April compared to June and October. These areas correspond to xeric, south-facing slopes, which, in the montane zone of the Front Range, mostly consist of open stands of ponderosa pine, compared to the more dense north-facing slopes (Veblen and Donnegan 2005). The more open forest floor may facilitate cougars using south-facing slopes as travel corridors, leading to greater transition rates. Similarly, Dickson et al. (2005) found that cougars used less rugged terrain than the surrounding area while traveling, while Knopff et al. (2014) found that cougars selected for south-facing slopes and areas with shrub habitat. The monthly difference in effect size for the response to heat loading may be related to seasonal changes in vegetation (shrub cover may be denser in June, reducing speed) or a product of unobserved weather patterns (e.g., more snow on north-facing slopes could lead to a greater tendency to use south-facing slopes as corridors). Similarly, we also observed slower movements at high elevations in April compared to June and October; high elevation areas of the Front Range may still contain snow in April, which could result in slower movement rates at high elevations. The time-varying directional response to elevation indicates that individuals are moving to higher elevations during the day, and then toward lower elevations at night. Blecha (2015) found that predation events, which typically occur at night, occurred at lower elevations, which may explain the temporal pattern we observed. The difference among seasons for the dawn motility response (slower [faster] at high [low] elevations in April, faster [slower] at high [low] elevations in October) might also be a response to unobserved fine-scale vegetation changes or dietary shifts.

The documented response of cougars to disturbed and developed landscapes varies in the literature, and is likely a function of the level of disturbance encountered. For example, Kertson et al. (2011) found no difference in cougar movement rates in wildland and residential areas throughout the day. However, Knopff et al. (2014) observed that cougars avoided developed landscapes, while also noting a temporal shift in usage of those areas, with cougars avoiding areas near development more during the day. On average, we observed a more negative relationship to distance from structures at dawn and dusk compared to mid-day and evening (i.e., individuals moved faster when closer to structures during dawn and dusk than mid-day and evening, when there was a slight positive relationship between speed and distance to structure), which could be explained by increased human activity caused by the start and end of the workday. In contrast, we observed orientation toward roads just after dawn in April and June and just before dawn in October, with orientation away from roads mid-day. However, the uncertainty was fairly large for the diel effect of distance to nearest roofed structure, despite subtle positive and negative shifts. These minor differences, and an overall lack of consistent response, could be explained by unmeasured spatial and temporal relationships, such as individual interactions and fine-scale human disturbance (e.g., recreational activities, noise, construction). For example, Wilmers et al. (2013) found that cougars showed stronger avoidance of more consistent sources of anthropogenic disruption, such as neighborhoods, than intermittent sources, such as low-traffic roads. We detected a faster average movement rate in areas with more development, which is consistent with work by Dickson et al. (2005) and Wang et al. (2017), who found that individuals expended more calories and moved further in developed areas.

Other studies have detected significant individual variation (Kertson et al. 2011; Wilmers et al. 2013), and Benson et al. (2016) and Wilmers et al. (2013) found that selection differed between males and females. We did not see consistent sex-specific responses to covariates, which could be due to the timing of the observations; for example, females may respond differently to males when breeding, but similarly at other times. In addition, males were underrepresented in our sample. Some of the unexplained individual variation could be due to the amount of anthropogenic landscape features each individual was likely to encounter in their movements Benson et al. (2016); Knopff et al. (2014), as opposed to the amount of development in the immediate vicinity during a given movement. Benson et al. (2016) also hypothesized that the amount of development in many studies of cougar habitat selection has been too low to cause cougar behavioral changes. We did not to detect a link between cougars that had reports of human conflict and response to development. Cougars have demonstrated different second- and third-order selection to roads in previous studies (Dickson and Beier 2002), therefore, individuals that become nuisance individuals may select for, or end up in, home ranges near human development, but do not respond differentially to areas closer to development within their home range (Linnell et al. 1999).

Many of the hypothesized movement drivers did not have a consistent statistically significant relationship with movement, while other studies on cougars in the same geographic area have found strong effects for landscape variables on cougar resource use while hunting (Blecha 2015). However, unlike RSFs and SSFs, the CTMC framework is measuring the effect of landscape variables on speed and directionality, not habitat preference. For example, cougars may select for areas with high mule deer use (Blecha 2015), but cougars may not alter their speed based on the amount of mule deer usage. In addition, the CTMC framework is based on a continuous-time movement path; although the model appropriately accounts for the uncertainty in the individual’s true location when it was not observed, it does introduce an additional source of variation that is not accounted for when using only the observed locations.

We propose that our findings regarding drivers of movement may have five potential biological causes. First, cougars are generalists, therefore, they are expected to demonstrate less habitat selection at the landscape scale than a habitat specialist would (Katnik and Wielgus 2005). Second, significant individual-variation within a 24-hour period would make determining a consistent population-level response difficult. Individual variation may be a function of the unmeasured internal state of the animal, such as breeding status, body condition, energetics, or interactions with other individuals. Future studies could focus on monitoring a range of individuals of known age, breeding status, and body condition. Third, it is possible that the risks and rewards present in the Front Range are not significant enough to cause a detectable response in behavior. Fourth, cougars may be responding to the landscape at a much finer scale than researchers are currently able to measure at such a large spatial extent. Finally, cougar movement may correspond to, or interact with, lower frequency environmental variation (e.g., weather patterns and food availability). Comparing behavior among years could be used to assess seasonal consistency in observed patterns; however, due to the difficulty of performing multiple comparisons of time-varying effects, a study of among year differences would likely need to focus on a particular season of interest.

The varying coefficient modeling framework, implemented in this study as a GAM, can reveal hidden process dynamics (Fan and Zhang 2008) and allows for complex nonlinear patterns that would be difficult to model in a traditional framework (e.g., Polansky and Robbins 2013). While we expanded each parameter in to the temporal space, one could make each covariate a function of another parameter, such as a different temporal predictor (e.g., time since kill) or another parameter in the model (e.g., distance to structure). Allowing the coefficients to vary in time (or another covariate space) can also improve the predictive ability of the model, as it did in our study. Conventional GLMs can mask time-varying responses to covariates (e.g., Cheng et al. 2009), because the response variable is aggregated over the time period of interest. Therefore, if the response of an individual switches between positive and negative (faster or slower movement rates), the estimated response will be approximately zero. Studies have found that cougars use a broader range of habitats for nocturnal movements than for daybed locations (Dickson et al. 2005) and demonstrate temporal variability in their response to anthropogenic landscape features (Knopff et al. 2014). Therefore, restricting analysis of locations to a particular temporal subset may not be indicative of all behavior (Comiskey et al. 2002). The CTMC framework represents an important step forward in detecting latent temporal patterns in animal movement and is especially useful when behavior is known to vary in time.

## MANAGEMENT IMPLICATIONS

We identified few significant population-level drivers of cougar movement, but we did identify several individuals that showed strong behavioral responses to landscape drivers. When making inference at a population-level, individual differences can negate one another (if some individuals respond positively and others negatively) or be overwhelmed by the non-response of other individuals. Therefore, individual variation can result in an inability to detect consistent patterns in behavior. For example, there was no detectable pattern in movement behavior that could be used for identifying individuals involved in conflict events (e.g., individuals involved in conflict move slower when near development). In addition, individuals moving within an established home range, such as in this study, may be acclimated (demonstrating minimal change in behavior) to the disturbances that they encounter during daily movements, making it difficult to extrapolate resident behavior to transient or dispersing individuals. The high degree of individual variation suggests that, if agencies want to minimize human-wildlife conflict, a “one size fits all” approach to cougar management and conflict abatement will likely be unsuccessful and management options should be varied and flexible. However, the strong speed and directional response to the repetitively used potential kill sites indicate that they are the driving force behind cougar movement behavior. Perhaps the most plausible and effective proactive mitigation of cougar conflict might be to identify potential kill sites and orient recreation (e.g., trails, camp sites) away from those areas. Communities and individual land owners could also use potential kill site information to inform land use within their property boundaries as a conflict mitigation strategy. Kill-site identification tools, such as those developed by Blecha (2015), could prove useful in this regard.

## ACKNOWLEDGEMENTS

Funding was provided by Colorado Parks and Wildlife (1304), the National Park Service (P12AC11099), Colorado Department of Transportation, NSF DMS 1614392, and NSF EEID 1414296. Data were provided by Colorado Parks and Wildlife (CPW ACUC #01-2008). Any use of trade, firm, or product names is for descriptive purposes only and does not imply endorsement by the U.S. Government.

## Associate Editor

**Summary for online Table of Contents**: Cougars in the wildland-urban interface of Colorado exhibit time-varying movement behavior in response to the landscape, primarily in response to the location of predation events. Not accounting for time-varying responses can mask drivers of movement.

## SUPPORTING INFORMATION

Additional supporting information may be found in the online version of this article at the publisher’s website.

## Appendix A

We spatially discretize a posterior predictive continuous path from the movement model to the resolution of the rasters of interest and decomposed into two elements: **c**, a state sequence consisting of the sequential grid cells (of *N* possible grid-cells) visited by the individual, and ***τ***, a vector of residence times that describe how long the individual spent in each grid cell. We describe the cell sequence in terms of the transition rates ***α*** where ***α*** _*ij*_ is a parameter controlling movement from cell *i* to cell *j* that can be a function of spatial covariates:

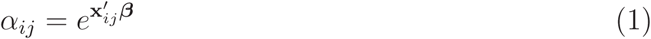

If we designate *t* as the *t*^*th*^ observation in the state-sequence (*t ∈ T*), then the residence time ***τ*** _*t*_ is exponentially distributed with a rate equal to the sum of all ***α*** _*ij*_ (the total transition rate):

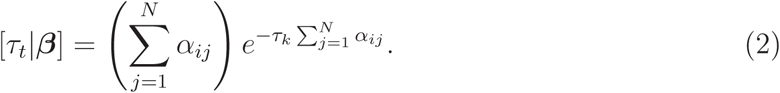

In the above notation, [***τ*** _*t*_|*β*] represents the probability distribution of the random variable ***τ*** _*t*_ given the parameters *β*; this notation will appear again. We assume that it is impossible to move directly to non-neighboring cells, and therefore ***α*** _*ij*_= 0 for all *j* except for the cells adjacent to cell *i*.

When an individual transitions to a neighboring cell, the probability of transitioning to cell *c* _*t*+1_ = *l* is

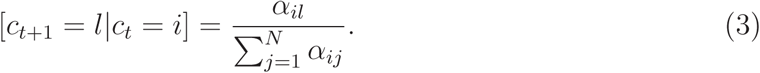

Assuming independence, the joint likelihood is the product of the transition probabilities and the residence times in the state sequence **c** is:

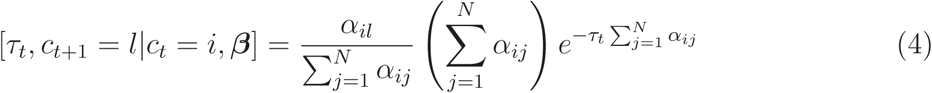

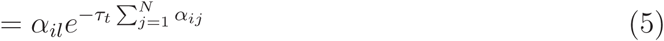

Using a latent variable representation, where

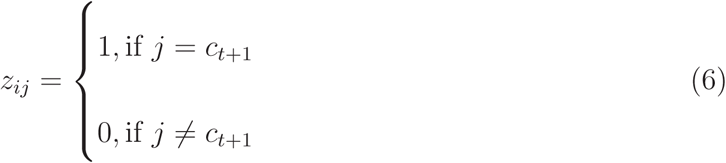

and

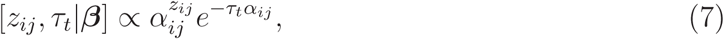

then the product of [*z* _*ctk*_, ***τ*** _*t*_|***β***] over all *N* is proportional to the likelihood of the observed transition:

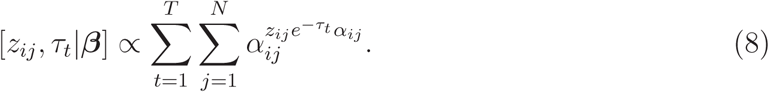

The above process is parameterized with a single realization from the movement model, however, we have failed to account for the uncertainty in the animal’s path. To avoid computational storage limitations, we use multiple imputation to account for the uncertainty in the path and make approximate posterior predictive inference on transition rates.

